# Development of novel cytoprotective small compounds inhibiting mitochondria-dependent apoptosis

**DOI:** 10.1101/2022.10.12.511987

**Authors:** Mieko Matsuyama, Joseph T. Ortega, Yuri Fedorov, Jonah Scott-McKean, Jeannie Muller-Greven, Matthias Buck, Drew Adams, Beata Jastrzebska, William Greenlee, Shigemi Matsuyama

**Affiliations:** Department of Ophthalmology and Visual Science; Department of Pharmacology and Cleveland Center for Membrane and Structural Biology; Department of Genetics and Genome Science; Department of Macromolecular Science and Engineering; Department of Physiology and Biophysics; Department of Pathology; Division of Hematology and Oncology. Department of Medicine; School of Medicine; School of Engineering; Case Western Reserve University Case Comprehensive Cancer Center; Harrington Discovery Institute Cleveland, OH

**Keywords:** Bax, Apoptosis, Cell Death, Retinal Diseases, Photoreceptor

## Abstract

We identified cyto-protective small molecules (CSMs) by a cell-based high-throughput screening of Bax inhibitors. Through a medicinal chemistry program, M109S was developed, which is orally bioactive and penetrates the blood-brain/retina barriers. M109S protected retinal cells in the mouse models of Stargardt disease and macular degeneration. M109S directly interacted with Bax and inhibited the conformational change and mitochondrial translocation of Bax. M109S inhibited ABT-737-induced apoptosis both in Bax-only and Bak-only MEFs. M109S also inhibited apoptosis induced by staurosporine (mouse embryonic fibroblasts), etoposide (Neuro2a cells), and obatoclax (ARPE19 cells). M109S is a novel small molecule protecting cells from mitochondria-dependent apoptosis both *in vitro* and *in vivo*. M109S has the potential to become a new research tool for studying cell death mechanisms and to develop therapeutics targeting mitochondria-dependent cell death pathway. (128words)

## Introduction

The Bcl-2 family of proteins is a group of evolutionarily conserved proteins regulating mitochondria-dependent apoptosis (Reviewed in (Singh et al., 2019)). The abnormal balance of these proteins induces apoptosis resistance of cancer cells and unwanted apoptosis of essential cells in the damaged tissues(Singh *et al*., 2019). Bax and Bak are the pro-apoptotic members of the Bcl-2 family of proteins playing a central role in the mitochondria-dependent programmed cell death (Wei et al., 2001). Since Bax/Bak double deficient cells are resistant to the stresses activating the mitochondria-dependent apoptosis pathway (Wei *et al*., 2001), Bax and Bak are ideal pharmacological targets to control the life and death of the cells. The pharmacological activators and inhibitors of Bax and Bak are expected to be effective in eliminating cancer cells and rescuing essential cells from unwanted death, respectively (Diepstraten et al., 2022; Jensen et al., 2019; Pogmore et al., 2021; Spitz and Gavathiotis, 2022; Walensky, 2019).

Previously, small compounds activating Bax/Bak-dependent apoptosis pathway have been developed, and some of these small compounds are used clinically as effective therapeutics eliminating cancer cells, for example, Venetoclax (Ashkenazi et al., 2017; Oltersdorf et al., 2005). However, clinically effective small compounds rescuing damaged cells from Bax/Bak-induced cell death have not yet been developed. Recent studies have reported the successful development of Bax-inhibiting compounds (pro-drugs) by identifying small molecules inhibiting the pore-forming activity (Niu et al., 2017) or the conformational change of Bax (Garner et al., 2019). In addition, Elthrompag, an FDA-approved drug for thrombocytopenia, was found to possess direct Bax binding and inhibiting activity (Spitz et al., 2021). These previous successes were achieved by using an artificially reconstituted *in vitro* assay system. The compounds identified by *in vitro* assay system will have the expected biochemical activity, however, there is no guarantee that they will have the expected efficacy in living cells without unexpected toxic effects. In this study, we utilized the cell-based functional screening system to develop novel CSMs that rescue cells from Bax-induced apoptosis. The chemical structures of CSMs, described here, are distinct from previously reported Bax inhibitors identified through the reconstituted *in vitro* system. Through medicinal chemistry efforts, we succeeded in developing M109S which has favorable pharmacokinetics in rodents. In this article, we report the chemical structures of CSMs and their activities protecting cells at the levels of cell culture and mouse retinal disease models.

## Results

### Development of cell-based Bax/Bak inhibitor screening system

We generated cell lines that undergo apoptosis by the expression of Bax or Bak in *bax^-/-^bak^-/-^*mouse embryonic fibroblasts (MEFs). In these cells, the Tet-ON promoter (Loew et al., 2010) is used for the induction of mCherry (red fluorescent protein), Bax, and Bak (Bax and Bak are expressed without mCherry-tag). Therefore, expression levels of Bax and Bak can be indirectly monitored by mCherry’s red fluorescence intensity. These cells were named iRFP (mCherry-inducible cells), iBax (Bax-inducible cells), or iBak (Bak-inducible cells) (Fig. 1A-C).

**Figure 1.**
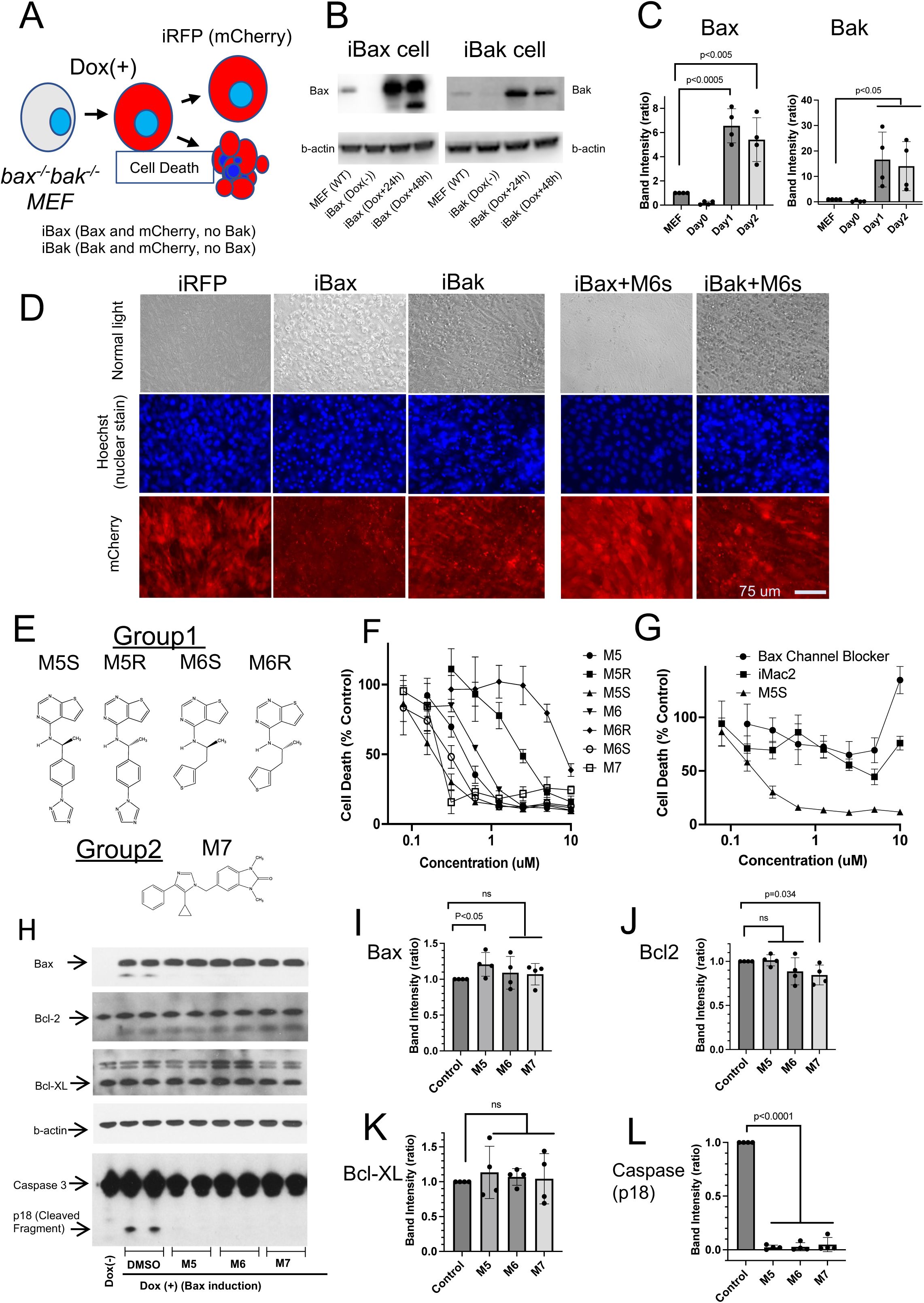
Development of Bax-or Bak-inducible system in *bax^-/-^bak^-/-^* MEFs and the screening of small compounds rescuing cells from Bax or Bak. A: Doxocyclin (Dox)-induced expression of Bax and Bak in MEFs. Dox-inducible Bax or Bak (no-tag) gene was introduced in *bax^-/-^bak^-/-^* MEFs by Lentivirus vectors. These cells were named iBax or iBak cells, respectively. The cells expressing only mCherry was named iRFP cells. B and C: Western blot analysis of the expression levels of Bax or Bak in wild-type (*wt*) MEFs, iBax, iBak cells are shown (B). The results of densitometric analyses are shown in C. The expression levels of Bax and Bak in *wt* MEFs were designated as 1 (actin levels were used as loading control). The error bars indicate SD. The result of the t-test statistical analysis is shown. Statistically significant difference (p<0.05) is indicated in the figure. D: Cell images of iRFP, iBax, and iBak. The Dox-inducible mCherry (red fluorescent protein) gene was introduced in *bax^-/-^bak^-/-^* to monitor Dox-inducible gene expression. Pictures were taken 48 h after Dox addition (100 ng/ml) to the medium. Images show cells under normal light, stained with Hoechst dye (nuclear staining), and expressing mCherry. Cell death (nuclear condensation and fragmentation), as well as apoptotic bodies (mCherry images), can be observed in these images. M6S is one of the hit compounds suppressing Bax-induced cell death. M6S (1 µM) rescued iBax cells, but not iBak cells. E: Chemical structures of hit compounds are shown. F: The dose-dependent inhibition of Bax-induced apoptosis in iBax cells by M5, M6, and M7 that were the best three representative hit compounds. The % of apoptosis was measured by the detection of apoptotic nuclei by Hoechst dye nuclear staining. The enantiomer S form of M5 and M6 (M5S and M6S) showed significantly stronger activities than the enantiomer R form (M5R and M6R). The original compounds in the library (M5 and M6) were a mixture of S and R forms with unknown ratio. G: The dose-dependent effects of M5S, iMac2, and Bax Channel Blocker suppressing cell death in iBax system is shown. M5S showed significantly stronger activity to inhibit cell death in iBax cells. H-L: Western blot analysis of Bcl-2 family proteins and Caspase-3 in iBax cells treated with M5, M6, and M7. iBax cells were incubated for 48 h with or without Dox (100 ng/ml) in the presence or absence of M5, M6, or M7. M5 and M6 are the original hit compounds that are a mixture of R and S forms. The results of the densitometric analysis are shown on the right as graphs. The expression level of each protein in the control group (DMSO) is designated as 1. (actin levels were used as loading control). All the compounds suppressed Caspase-3 cleavage without major changes in the expression levels of Bax, Bcl-2, and Bcl-XL. The result of the t-test statistical analysis is shown in each graph. “ns”: No statistically significant difference was detected (p>0.05).

Doxycycline (Dox), a tetracycline analog, was used to activate the Tet-ON system, and the expression of Bax and Bak was detected by Western blot at 6 h after the induction (Fig. 1B and C). Forty-eight hours after the induction, many apoptotic cells can be observed by microscopic analysis (Fig. 1D). To detect apoptotic nuclei, cells were stained with Hoechst dye (Fig. 1D). These iBax cells were used to identify the candidates of CSMs. A 50,000 small compound library was used to identify hit compounds by using a cell image analysis system (Operetta system, Parkin-Elmer). The hits were selected by the following two criteria; 1) Decreasing the percentages of dead cells detected (nuclear fragmentation and condensation), and 2) Maintaining mCherry expression (fluorescence intensity) above 80% of the control so that compounds inhibiting the Tet-ON system could be excluded. We then selected the top 50 compounds that satisfied these criteria. Based on similarities in the chemical structure, the compounds were divided into 2 groups. The best three small molecules rescuing iBax cells were M5, M6, and M7 (Fig. 1E). M5 and M6 (that have two enantiomers R and S) were assigned to Group 1, and M7 to Group 2. Fig. 1D shows an example of M6S protecting iBax cells from Bax-induced apoptosis while maintaining mCherry expression. These Bax-inhibiting hits were then tested in iBak cells. However, none of these compounds were able to inhibit cell death induced by Bak in iBak cells (Fig. 1D, example images of iBax and iBaK cells incubated with M6S 1 µM are shown). These results suggest that CSMs can rescue cells from the apoptotic processes induced by the overexpression of Bax but not Bak. The dose-response effects of M5, M5R, M5S, M6, M6R, M6S, and M7 inhibiting Bax-induced apoptosis in iBax cells are shown in Fig. 1F. The S-enantiomer showed a significantly higher cell death-inhibiting activity than the R-form in both M5 and M6. Therefore, we focused on the S-enantiomer to perform Hit-to-Lead optimization in Group 1. To be noted, the cell death inhibition activity of M5S is significantly higher than the activities of Bax Channel Blocker(Bombrun et al., 2003) and iMac2 (Bombrun *et al*., 2003; Peixoto et al., 2009)(Fig.1G). Western blot analyses showed that these compounds did not decrease Bax expression (Fig. 1H and I) but rather suppressed Caspase-3 activation (cleavage) (Fig. 1H and L). The expression levels of endogenous Bax inhibitors such as Bcl-2 and Bcl-XL did not show major changes (Fig. 1J and K).

### Hit-to-Lead optimization

A medicinal chemistry program was carried out to expand the two lead series and optimize iBax cell potency and Bax binding (explained later in Fig. 5E and G). A total of 200 new inhibitors were designed and synthesized. Among them, M41S (Series 1) and M11 (Series 2) were the best apoptosis inhibitors in iBax cells (Fig. 2A-C). The EC_50_ of M41S and M11 for the inhibition of caspase activation in iBax cells were 3.8 nM (Fig. 2D) and 145 nM (Fig. 2C), respectively. In comparison with the original hit compounds of M5 (S (EC_50_=124 nM) and R (EC_50_=1,450 nM)), and M7 (EC_50_=553 nM), apoptosis inhibiting activities of M41S and M11 were significantly improved (Fig. 2B-D). Further optimization of the lead compounds (M41S) led to the identification of the M109S lead compound protecting iBax cells with the EC_50_ of 23.4 nM (Fig. 2A and D). M109S exhibits drug-like profiles: Molecular Weight is 375, cLogP=4.0, kinetic solubility is 0.05 uM in pH7.4 (clear suspension/solution can be prepared in 15% beta-cyclodextrin solution at 2 mg M109S/ml), Caco-2 cell permeability is 11.7 x 10^-6^ cm/sec and the efflux ratio is 1.3, microsomal stability (half-life (t ½)) is 46 min (human) or 11 min (mouse), plasma protein binding is 99.97% (human) or 99.71 (mouse). The detail synthesis protocols and pathways for M109S has been published (Matsuyama, 2020). As shown in Fig.2D, the apoptosis inhibition activity of M41S and M109S in iBax cells is significantly higher than the activity of Elthompag, a thrombopoietin receptor analog, which is recently reported to have Bax inhibitor activity (Spitz *et al*., 2021).

**Figure 2.**
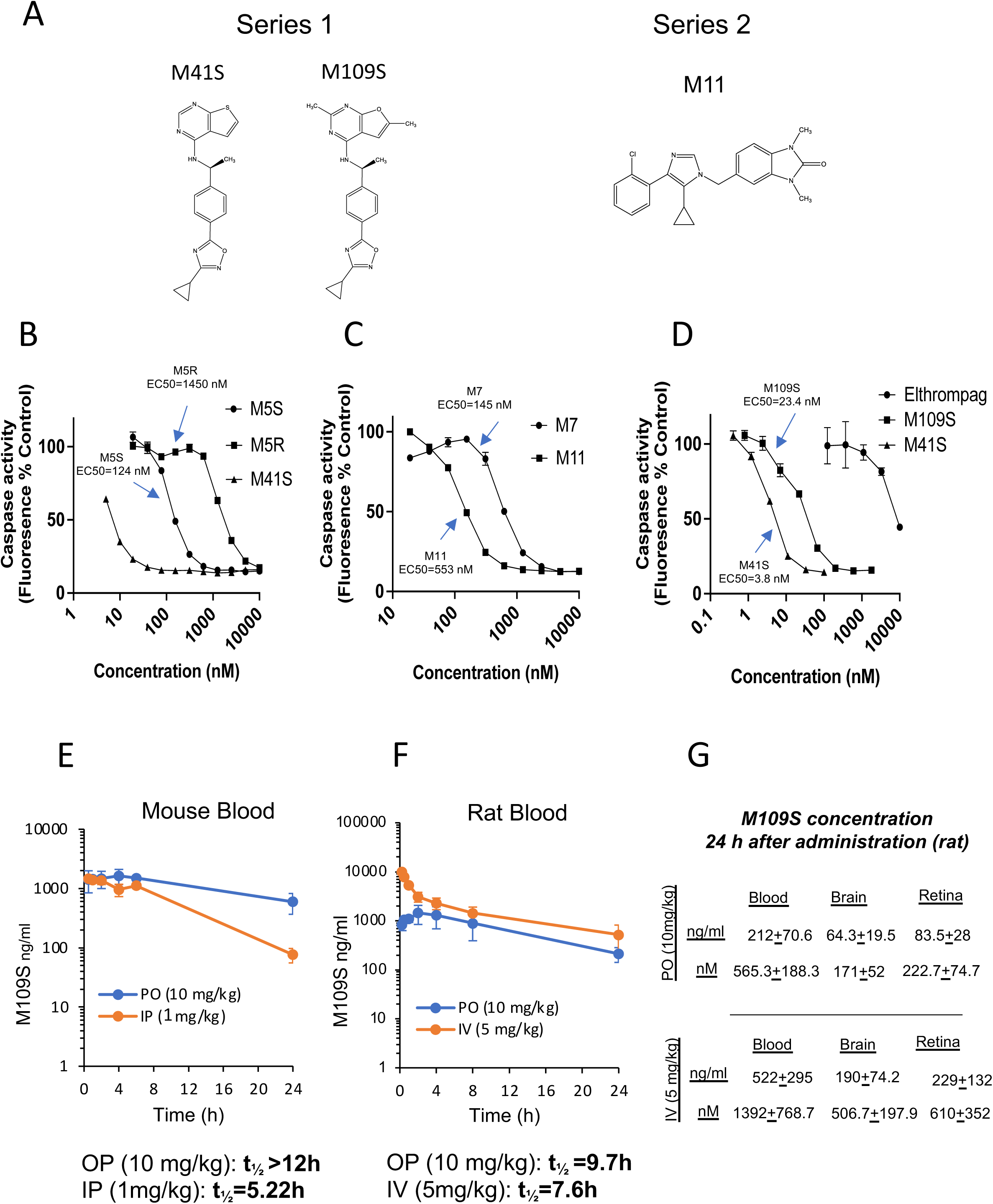
Hit-to-Lead optimization of the novel <u>c</u>ytoprotective <u>s</u>mall <u>molecules</u> (CSMs). A: Chemical structures of newly designed and synthesized lead compounds. The most effective cell death inhibitor in cell culture, M41S, was developed based on the common chemical structure of M5S and M6S (Series 1). The orally bioactive CSM, M109S, was developed from M41S. M11 was the most effective cell death inhibitor among the inhibitors designed from M7 (Series 2). B-D: Caspase activity of iBax cells treated with CSMs. Bax-mediated apoptosis was induced by Dox addition to the culture medium in iBax cells in the presence of various concentrations of CSMs as indicated in the graphs. The improvement of apoptosis inhibition through Hit-to-Lead optimization is shown in B (M41S vs M5S and M5R) and C (M11 vs M7). D shows the comparison of M41S (the best cell death inhibitor in cell culture), M109S (orally bioavailable CSM designed from M41S), and Elthrmpag (previously reported Bax inhibitor). E-G: Pharmacokinetic analysis of M109S in rodents. M109S was administered to mice and rats by intraperitoneal injection (IP, 1 mg/kg), intravenous injection (IV, 5 mg/kg), or oral gavage (PO, 10 mg/kg). The concentration of M109S was measured by Mass Spectrometry analysis. G: M109S concentrations in the blood, brain, and retina 24 h after the administration (oral gavage (PO) or intravenous injection (IV)) in rats.

### M109 is an orally bioactive cell death inhibitor penetrating blood-brain/retina-barrier

The pharmacokinetic (PK) analysis of M109S was performed in both mice and rats by oral gavage (PO) and either by intraperitoneal (IP) or intravenous injection (IV) (Fig. 2E-G). In mice, M109S reached 1.0 μg/ml (2.6 µM) plasma concentration within 30 min from administration, and it remained at 596 ± 134 ng/ml (1.6 ± 0.36 µM) 24 h after the oral gavage administration (Fig. 2E). Similar results were obtained in rats (Fig. 2F). At 24 h after the oral gavage administration, the level of M109S in the plasma was 565.3 ± 188.3 nM in rats. Moreover, M109S crossed the blood-brain and blood-retina barriers. The level of M109S in the rat retina and brain reached 171.0 ± 52.0 nM and 222.7 ± 74.7 nM, respectively, 24 h after its oral administration (Fig. 2F and G). Notably, these concentrations are higher than the EC_50_ concentration (23.4 nM) of M109S protecting iBax cells (Fig.2D). The half-life (t_1/2_) of M109S (10 mg/kg body weight (b.w.) oral administration) in the blood was longer than 12 h (in mice) or 9.7 h (in rats) (Fig. 2E and F).

### M109S protected the retina from the bright light-induced photoreceptor death

Previous studies showed that Bax-mediated apoptosis plays an essential role in bright light-induced retinal degeneration in *Abca4^−/−^Rdh8^−/−^* mice(Sawada et al., 2014). *Abca4^−/−^Rdh8^−/−^* mice are mouse models of human Stargardt disease, a juvenile age-related macular degeneration (AMD) that is susceptible to acute light insult and long-term retinal degeneration(Maeda et al., 2009) (Sawada *et al*., 2014) (Ortega et al., 2021). The retinal photoreceptor cells degenerate within 7 days in these mice after illumination with bright light due to apoptosis. An oral administration of these mice with a total of four doses of M109S at 10 mg/kg b.w. within a scheme of two doses before light exposure (24 h and 1h), followed by two doses after light exposure (24 h and 48 h), produced protection of photoreceptor cells from death (Fig. 3A-H). The apoptotic processes occurring in the photoreceptor cells upon bright light injury trigger the activation of an immune response, which correlates with an increased number of autofluorescence (AF) spots in the retina. The AF spots that can be detected with Scanning Laser Ophthalmoscopy (SLO) imaging represent the activated resident microglia and peripheral immune cells that migrate to the retina to clear dying photoreceptors. The increased number of AF spots was detected only in the vehicle-treated mice. However, in mice treated with M109S at 10 mg/kg b.w., the number of AF spots was similar to that detected in the dark-adapted mice (Fig. 3A, B, and S2). The loss of photoreceptors in the vehicle-treated mice was evidenced by thinning of the outer nuclear cell layer (ONL) detected by the optical coherence tomography (OCT), while the ONL in mice treated with M109S at 10 mg/kg b.w closely reassembled the ONL in unexposed mice (Fig. 3C and D). While lower doses were less effective (Fig. 3D and S3). The thickness of the ONL was confirmed by histological examination of the H&E-stained retinal sections (Fig. 3E and G). Immunohistochemical staining of rod and cone cells revealed healthy photoreceptors in mice pretreated with M109S prior to bright light injury (Fig. 3F). The protection of the photoreceptor cells was confirmed in non-pigmented WT Balb/cJ mice sensitive to light-induced retinal degeneration (Fig. 3H and S4). While exposure to bright light triggered deterioration of photoreceptor cells in the vehicle-treated mice, oral administration of M109S 24 h and 1 h prior to illumination followed by additional treatment 24 h and 48 h after light exposure protected these mouse retinas. Altogether, these results indicate that M109S protects retinal cells from bright light-induced death, and thus prevents retina degeneration triggered by illumination.

**Figure 3.**
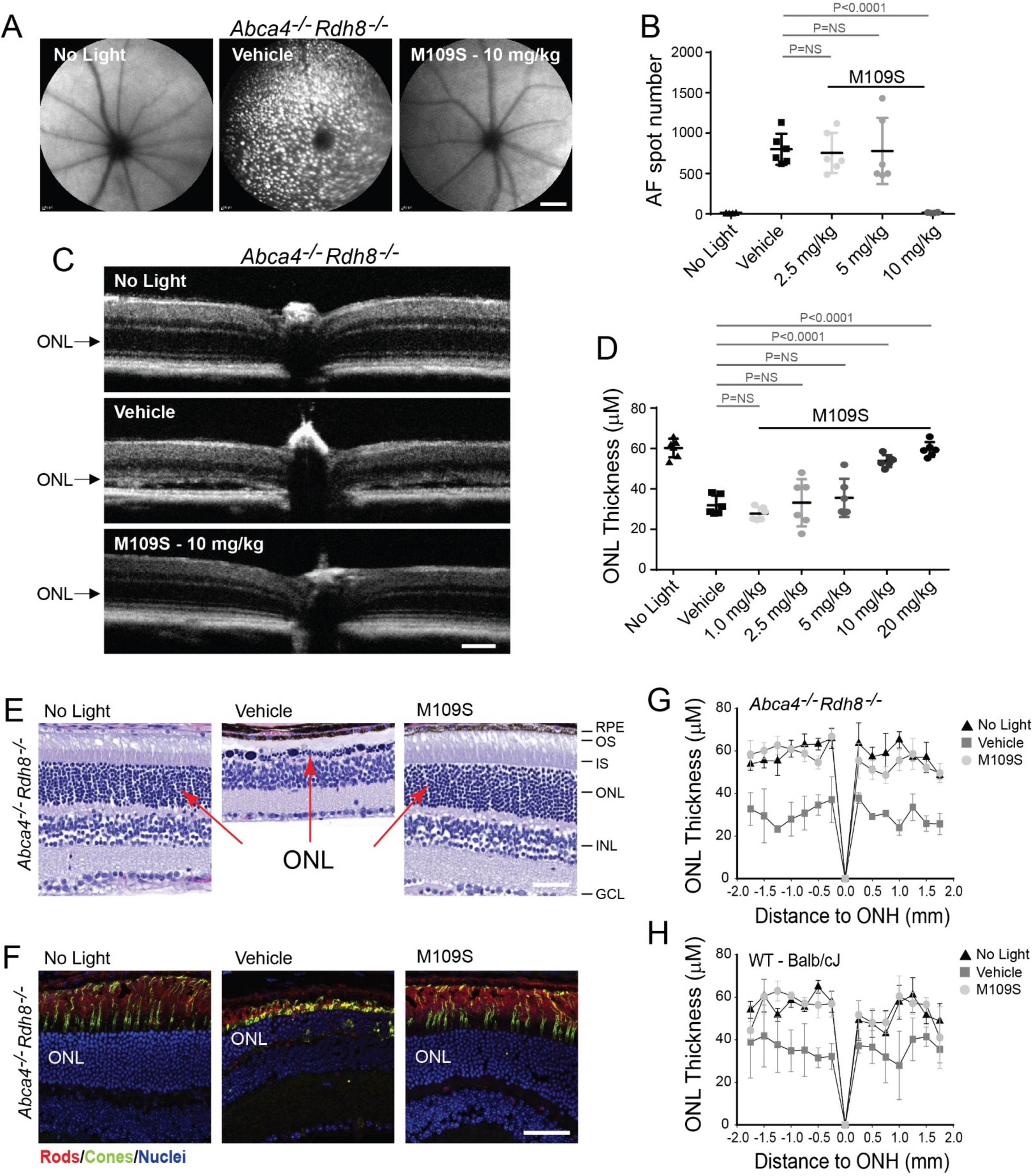
M109S protects the retina from bright light-induced injury in *Abc4^-/-^Rdh8^-/-^* mice and Balb/cJ mice. A and B: A, The *in vivo* whole fundus imaging from *Abca4^-/-^Rdh8^-/-^* mice taken with scanning laser ophthalmoscopy (SLO) one week after exposure to bright light. Autofluorescence spots (AF) representing macrophages and microglial cells activated by light injury to clear dying photoreceptors were detected in only vehicle-treated mice, but not in M109S-treated mice (10 mg/kg/b.w.). Scale bar, 1 mm. B, Quantification of the AF. M109s at 10 mg/kg/b.w showed statistically significant protection against light-induced photoreceptor cell death. Statistical analysis was performed with the two-way ANOVA and post hoc Turkey tests. The values of p<0.05 were considered statistically significant. No statistically significant changes were labeled as “ns”. C and D: C, The Spectral Domain Optical Coherence Tomography (SD-OCT) images of the retina from *Abca4^-/-^Rdh8^-/-^* mice taken one week after illumination. Scale bar, 100 µm. D, The measurement and statistical analysis of the ONL thickness in SD-OCT images. The exposure to bright light-induced the degeneration of photoreceptors, which caused the shortening of the outer nuclear layer (ONL) in retinas in vehicle-treated control mice, but not in the M109S-treated group. M109S showed a dose-dependent effect protecting the photoreceptor cells from light-induced death. Statistical analysis was performed with the two-way ANOVA and post hoc Turkey tests. The values of p<0.05 were considered statistically significant. No statistically significant changes were labeled as “ns”. E, The hematoxylin and eosin (H&E) staining of the retina sections of *Abca4^-/-^Rdh8^-/-^*mice. M109S treatment mitigated degenerative processes in the retina. The thickness of the ONL in mice treated with M109S at 10 mg/kg b.w. closely resembled the thickness of the retina in dark-adapted mice. Scale bar, 50 µm. F, Immunohistochemical analysis of the retina of *Abca4^-/-^Rdh8^-/-^*mice. The cryosections were stained with an anti-rhodopsin antibody recognizing the C-terminus (1D4 antibody) to detect rod photoreceptors (red) and with peanut agglutinin to detect cone photoreceptors (green). Nuclei were stained with DAPI (blue). M109S treatment prevented the degeneration of both rod and cone photoreceptors. Scale bar, 50 µm. G and H: The measurement and statistical analysis of the thickness of the ONL in the H&E-stained sections of the retina of *Abca4^-/-^Rdh8^-/-^* mice (G) and Balb/cJ mice (H). The treatment with M109S (10 mg/kg/b.w.) treatment prevented the thickening of the ONL caused by the bright light injury in both mouse strains.

### Evaluation of cell death inhibition activities of M109S in cultured cells

Next, we evaluated the effectiveness of M109S inhibition of cell death pharmacologically induced in different cell types. ABT-737 is known to activate Bax/Bak-dependent apoptosis pathway (Certo et al., 2006) (Vogler et al., 2009). The effects of M109S against ABT-737-induced apoptosis were examined in MEF cells of wild type (*wt*) (Fig. 4A), *bax^+/+^bak^-/-^*(Bax only) (Fig. 4B), and *bax^-/-^bak^+/+^* (Bak only) (Fig. 4C). M109S showed a dose-dependent suppression of caspase activation in all three types of MEFs, suggesting that M109S can inhibit apoptosis induced by Bax as well as Bak which are expressed at endogenous levels. Staurosporine (STS), a pan-kinase inhibitor, is often used as a cytotoxic chemical activating Bax/Bak-dependent mitochondria-induced apoptosis (Wei *et al*., 2001). M109S inhibited apoptosis induced by STS in MEFs (Fig. 4D). Etoposide, a topoisomerase inhibitor, is known to activate DNA damage-induced cell death involving Bax/Bak-mediated apoptosis (Han et al., 2019; Tu et al., 2009). M109S also significantly suppressed apoptosis induced by etoposide in Neruo2a cells (Fig. 4E). However, in HeLa cells, M109S showed only slight inhibition (most of the effects were not statistically significant) of etoposide-induced apoptosis (Fig. 4F and Fig. S4), indicating that M109S inhibits etoposide-induced apoptosis in a cell type-dependent manner. Obatoclax is known to induce both Bax/Bak-dependent and independent apoptosis in certain types of cancer cells (Vogler *et al*., 2009). M109S inhibited obatoclax-induced apoptosis in ARPE19 cells (retina cell derived cancer cells). M109S suppressed caspase-3 activation without a significant impact on Bax levels of obatoclax-treated ARPE19 cells (Fig. 4 G-I).

**Figure 4.**
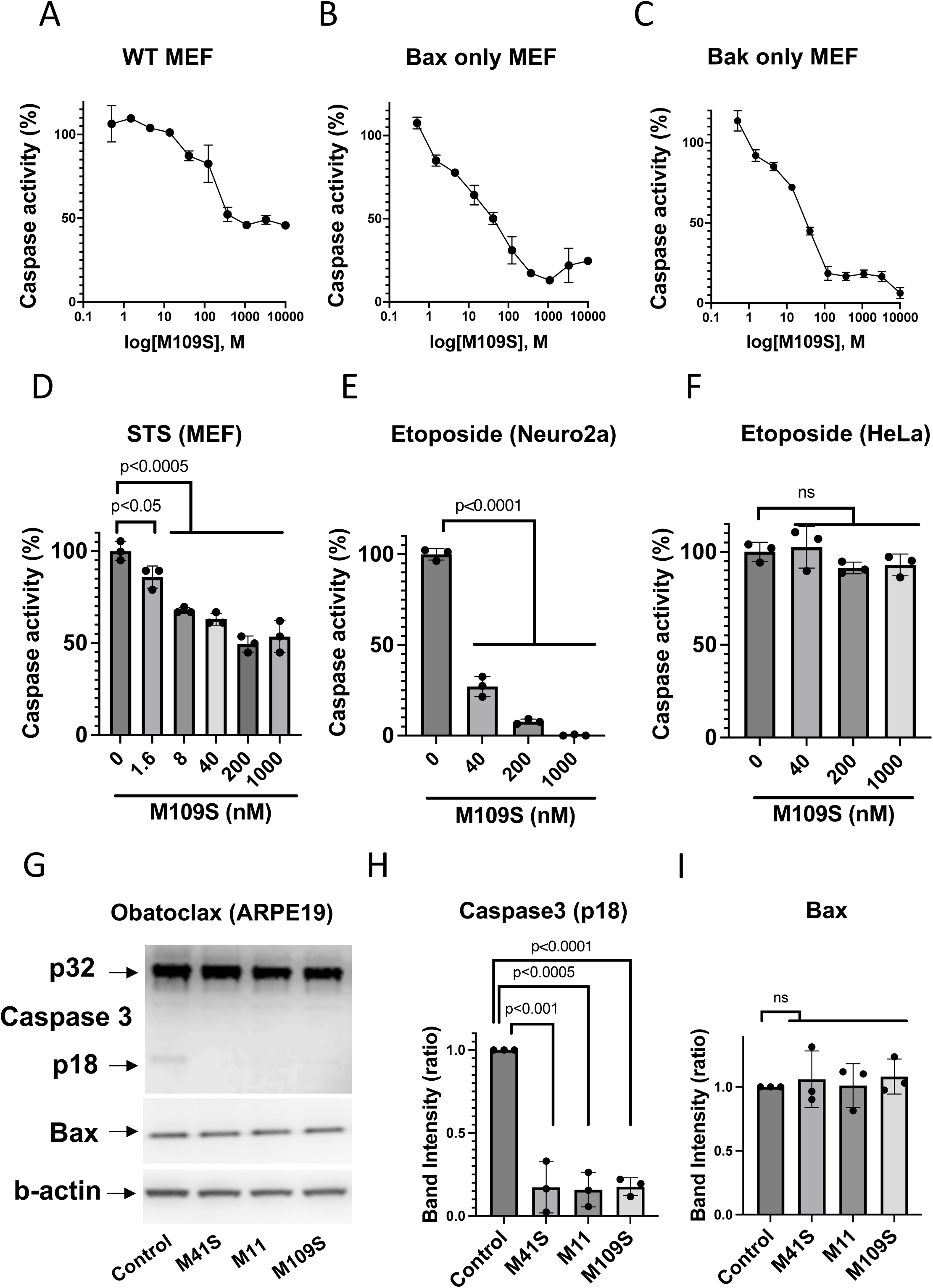
Effects of M109S on drug-induced apoptosis in cell culture. A-C: M109S suppressed apoptosis induced by ABT-737 wild type (A), Bax only (*bax^+/+^bak^-/-^*) (B), and Bak only (*bax^-/-^bak^+/+^*) (C) MEFs. MEFs were cultured with ABT-737 (1 µM) with various doses of M109S for 24 h (WT and Bax only) or 48 h (Bak only). Caspase activitiy of vehicle control (ABT-737 and vehicle) was designated as 100%. D: M109S suppressed staurosporine (STS)-induced apoptosis in MEFs. MEFs were treated with STS (1 µM) for 4 h in the presence of various concentrations as indicated in the graph. M109S suppressed STS-induced caspase activation in a -dose-dependent manner. The result of the Student’s t-test statistical analysis is shown in each graph. “ns”: No statistically significant difference was detected (p>0.05). E: M109S inhibited etoposide-induced apoptosis in Neuro2a cells. Neuro2a cells were treated with etoposide (12.5 µM) for 24 h. Then, the medium was changed, and cells were incubated with various concentrations of M109S for an additional 24 h without etoposide. M109S suppressed etoposide-induced caspase activation. The result of the Student’s t-test statistical analysis is shown in each graph. “ns”: No statistically significant difference was detected (p>0.05). F: M109S was not able to inhibit etoposide-induced apoptosis in HeLa cells. HeLa cells were treated with Etoposide (12.5 µM) for 24 hrs. Then, the medium was changed, and cells were incubated with various concentrations of M109S for an additional 24 h without Etoposide. M109S did not show significant inhibition of Etoposide-induced caspase activation in HeLa cells. The graph shows the results of 12.5 µM Etoposide. The result of the Student’s t-test statistical analysis is shown in each graph. “ns”: No statistically significant difference was detected (p>0.05). G-I: M109S inhibited Obatoclax-induced apoptosis in ARPE19 cells. AREP19 cells were incubated with Obatoclax (100 nM) in the presence of 500 nM CSMs (M41S, M11, and M109S) for 24 h. Then, cells were collected and Western analyses of Caspase-3 and Bax were performed. CSMs suppressed obatoclax-induced Caspase activation (p18 fragment production by the cleavage of Caspase-3) (G). Bax expression was not affected by CSMs (I) (actin levels were used as loading control). The result of the Student’s t-test analysis is shown in each graph. “ns”: No statistically significant difference was detected (p>0.05).

### M109S suppressed the conformation change (N-terminal exposure) and the mitochondrial translocation of Bax

The exposure of the N-terminus of Bax occurs as the early step of Bax-induced apoptosis (Hsu and Youle, 1998; Nechushtan et al., 1999). The levels of this conformational change can be examined by immunoprecipitation of Bax by antibodies recognizing the N-terminus of Bax (Hsu and Youle, 1998). As shown in Fig. 5A and B, CSMs (M41S, M11, and M109S) suppressed the conformational change of Bax. Bax is known to translocate from the cytosol to the mitochondria after the conformational change, which could be detected as higher density puncta in the DMSO-treated control cells labeled with anti-Bax antibody (Fig. 5 C and D). (Nechushtan *et al*., 1999). CSMs (M41S, M11, and M109S) suppressed mitochondrial translocation of Bax in iBax cells (Fig. 5 C and D).

**Figure 5.**
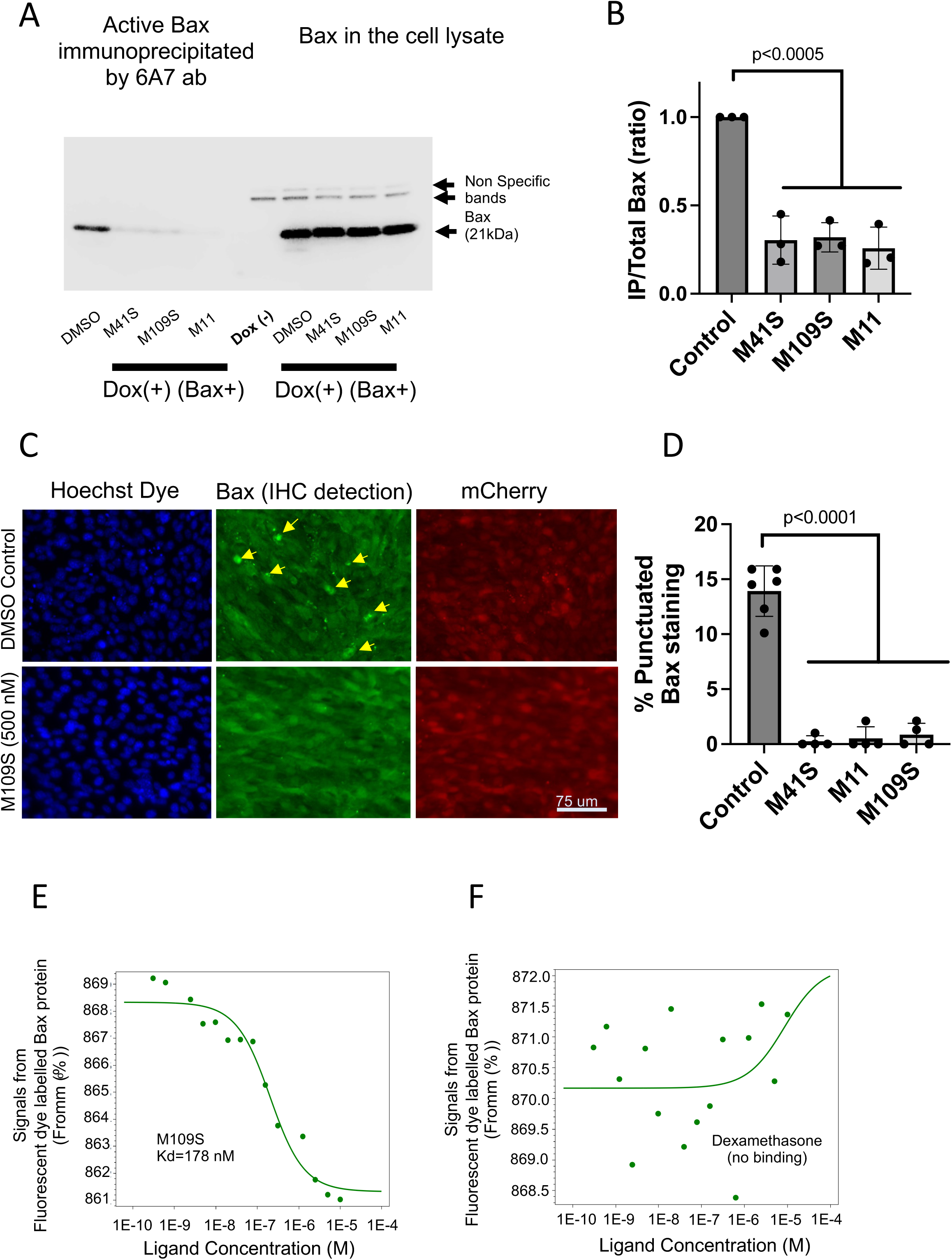
M109S inhibits Bax activation. A and B: CSMs suppressed conformation change of Bax. The N-terminus exposure of Bax occurs at the early step of Bax activation. Activated (conformationally changed) Bax can be immunoprecipitated by the 6A7 monoclonal antibody recognizing the N-terminus of Bax. iBax cells were treated with Dox for 48 hrs in the presence or absence of CSMs (M41S (50nM), M109S (500nM), or M11 (500nM)). Cells were lysed with CHAPS buffer and the lysates were subjected to immunoprecipitation (ip). C: Western blot of immunoprecipitated Bax by 6A7 antibody and total Bax in the cell lysates are shown. D: The ratio of the Western blot intensity of immunoprecipitated Bax (IP) and total Bax (total) is shown. CSMs significantly suppressed the amount of immunoprecipitated Bax without a significant change in the total Bax expression. The result of the Student’s t-test statistical analysis is shown in each graph. “ns”: No statistically significant difference was detected (p>0.05). C and D: CSMs suppressed mitochondrial translocation of Bax. iBax cells were fixed at 36 h after Dox treatment. Bax was detected by an anti-Bax polyclonal antibody (green fluorescence). The activation of the Dox-inducible promoter was confirmed by the detection of mCherry by red fluorescence. In vehicle (DMSO) control, punctuated Bax staining patterns associated with mitochondrial translocation were detected (indicated by arrows). The frequency of the punctuated staining was significantly reduced by CSMs (M41S, M11, and M109S). D: Quantitative analysis of Fig.5C is shown. The result of the Students’ t-test statistical analysis is shown in graph (B). “ns”: No statistically significant difference was detected (p>0.05). E and F: CSMs bind purified recombinant Bax proteins. The interactions of CSMs and Bax were examined by microscale thermal shift (MST) assay using His-tagged Bax. Panel E shows a representative dose-dependent Bax binding signal curve of M109S. A similar binding assay was performed multiple times, and the average of Kd was 153.75+55.8 nM (n=4). Panel F shows a negative control experiment. The binding of Bax and Dexamethasone was not detected.

### M019S directly binds to recombinant Bax protein

We examined whether CSMs (M109S and M41S) directly bind Bax using a recombinant Bax protein (Fig. 5E and S6). Binding was examined by the microscale thermophoresis (MST) assay (Wienken et al., 2010). M109S showed Bax binding activity with the average Kd of 153.75±55.8 nM calculated from four independent MST assay results. Fig.5E shows an example of MST assay detecting the biding of M109S and Bax (Kd=178nM). Dexamethasone is a steroid hormone analog that has been shown to inhibit fibroblast cell death (Amsterdam et al., 2002; Gascoyne et al., 2003), but it is not expected to bind Bax. Dexamethasone was used as a negative control, and the binding of Dexamethasone and Bax was not detected (Fig. 5F).

### M109S decreases mitochondrial oxygen consumption and reactive oxygen species whereas M109S increases glycolysis

We noticed that the color of the culture medium of M109S-treated cells turned from red to orange a little bit faster than that of vehicle control, suggesting that the medium acidification was stimulated by M109S. Since lactate accumulation as the result of glycolysis is the common reason for medium acidification in cell culture, we speculated that M109S has an activity influencing cellular metabolism. To determine the effects of M109S on cellular metabolism, oxygen consumption rate (OCR) and extracellular acidification rate (ECAR) were measured using the Seahorse instrument. OCR and ECAR are the indicators of mitochondrial oxidative phosphorylation (OXPHOS) and glycolysis, respectively. At 1 μM concentration in MEF cell culture, M109S decreased maximal OCR and increased ECAR (Fig. 6A, C, E, and H). Interestingly, similar effects were observed in *bax^-/-^bak^-/-^*(Bax/Bak double knockout (DKO)) MEFs (Fig. 6B, D, F, and H), indicating that this activity on metabolism is independent of Bax and Bak. Previous studies showed that OXPHOS inhibitors have neuroprotective activities by decreasing reactive oxygen species (ROS) generation from mitochondria (Procaccio et al., 2014) (Cunnane et al., 2020; Gao et al., 2021; Stojakovic et al., 2021). Therefore, the effects of M109S on ROS were examined. As shown in Fig. 6I and J, M109S decreased ROS levels both in *wt* and *bax^-/-^bak^-/-^* DKO MEFs. N-acetyl cysteine (NAC) is known to reduce ROS levels in cells (Ezerina et al., 2018). The effects of M109S (0.1-1 µM) on the ROS level in cultured MEFs were very similar to the effects of NAC (10-20 µM) (Fig.6I). Since the changes in the metabolism can impact the speed of cell division, we examined the effects of M109S on population doublings of cultured cells. M109S slightly decreased the speed of the population-doubling time of cultured MEFs (Fig. 6K and L). The effect was more evident in *bax^-/-^bak^-/-^* MEFs than in *wt* MEFs. Probably M109S’s effects on cell division speed become more detectable in *bax^-/-^bak^-/-^* MEFs because the population doubling speed of *bax^-/-^bak^-/-^* MEFs is faster than that of *wt* MEFs. Altogether, these results indicate that independently of Bax inhibiting effect, M109S also decreases the rate of oxygen consumption, which diminishes the levels of ROS in the cell. These different activities of M109S have likely additive protective effects against mitochondria-dependent cell death pathways.

**Figure 6.**
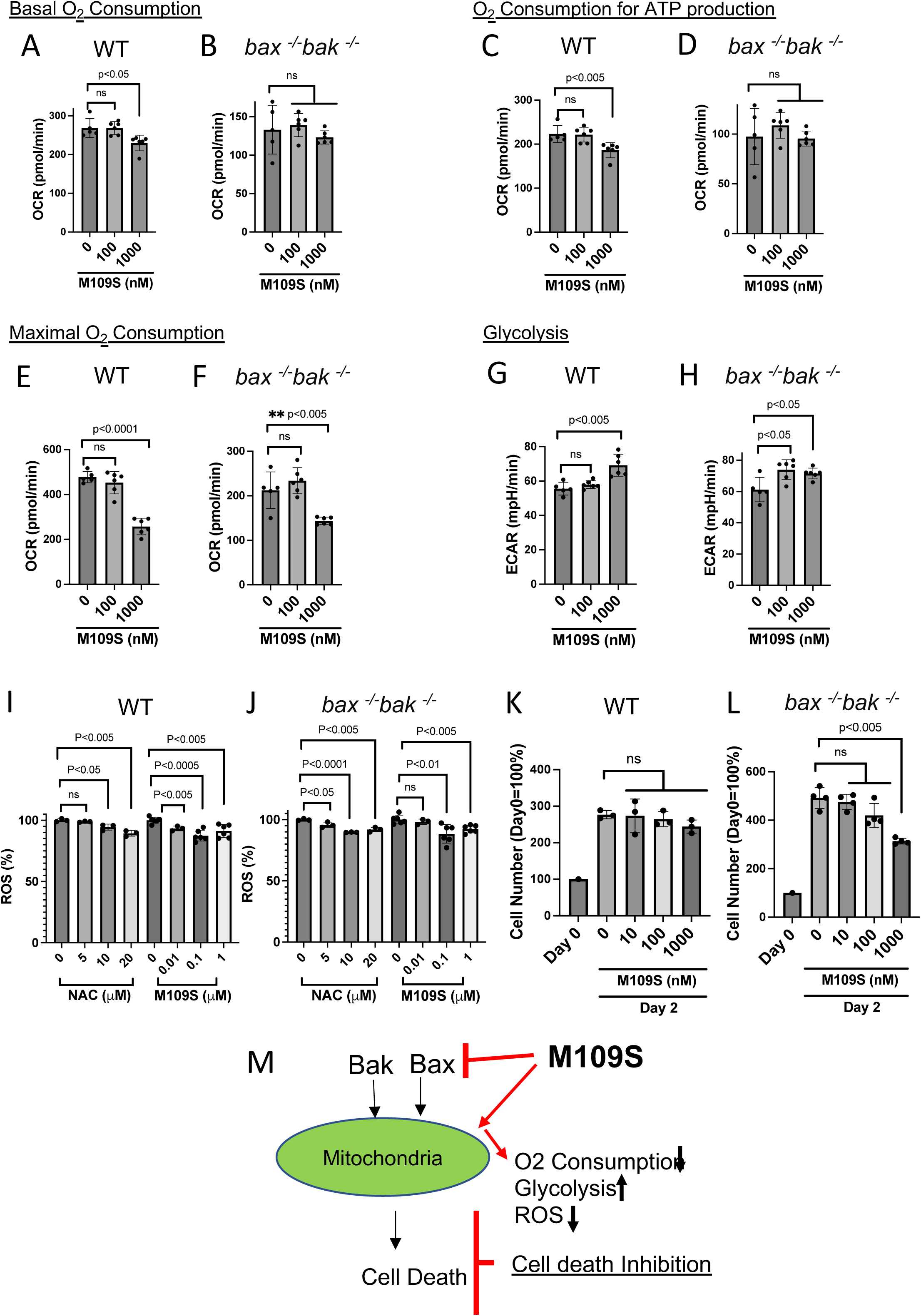
Effects of M109s on cellular metabolism. A-H: M109S decreased the maximum oxygen consumption rate (OC<u>R</u>) and increased the extracellular acidification rate (ECAR) both in *wt* and *bax^-/-^bak^-/-^* MEFs. OCR and ECAR represent the activities of mitochondrial oxidative phosphorylation (OXPHOS) and glycolysis, respectively. The effect of M109S on metabolism was analyzed using the Seahorse mitochondria stress kit as described in Method section. MEFs (*wt* and *bax^-/-^bak^-/-^*) were incubated with or without M109s for 4 h before Seahorse analysis. M109S was also added to the culture medium during the Seahorse analysis procedure (approximately 3 h). A-F: The levels of OCR of the basal (A, B), ATP production (C, D), and maximum oxygen consumption (E, F), are shown. G and H: The basal ECAR is shown. M109S decreased maximal O_2_ consumption and increased glycolysis in *wt* and *bax^-/-^bak^-/-^* MEFs. The result of the Student’s t-test statistical analysis is shown in each graph. “ns”: No statistically significant difference was detected (p>0.05). I and J: M109S decreases ROS levels in cell culture. ROS levels were measured by ROS-Glo-H_2_O_2_ assay kit (Promega). N-acetyl cysteine (NAC) or M109S was added to the culture medium for a total of 8 h (4 hrs before ROS substrate addition and 4 h during the incubation with ROS substrate). M109S addition decreased ROS levels both in *wt* and *bax^-/-^bak^-/-^* MEFs. The result of Studen’s t-test statistical analysis is shown in each graph. “ns”: No statistically significant difference was detected (p>0.05). K and L: A high dose of M109S (1 µM) slowed the cell division speed both in *wt* and *bax^-/-^bak^-/-^*MEFs. Forty thousand cells were plated in each well of 12 well plate (1 ml/well) on Day 0. The next day (24 h later), M109S was added to the medium, and cells were cultured for an additional 24 h, and the numbers of cells were counted. The cell number at Day 0 is designated 100%, and the ratio of cell number is shown. The result of the Student’s t-test statistical analysis is shown in each graph. “ns”: No statistically significant difference was detected (p>0.05). M: Schematic explanation of the mechanism of action of CSMs.

## Discussion

We developed CSMs by utilizing a cell-based functional assay system that was developed to identify Bax inhibitors. The chemical structures of these CSMs are distinct from previously reported Bax inhibitors (Niu *et al*., 2017) (Amgalan et al., 2020; Garner *et al*., 2019; Spitz *et al*., 2021) that were identified through *in vitro* Bax binding and pore-forming assays. Through a medicinal chemistry program, M109S was developed as a lead compound which possesses cell protecting activities both *in vitro* and *in vivo*. M109SS suppressed ABT-737-induced apoptosis in both Bax-only and Bak-only MEFs (Fig. 4C). These observations suggest that M109S can inhibit apoptosis induced by Bax and Bak expressed at endogenous levels in the cell. M109S also inhibited apoptosis in STS-treated MEFs, obatoclax treated ARPE19 cells, and etoposide-treated Neuro2a cells (Fig. 4 A-I). However, M109S did not inhibit etoposide-induced apoptosis in HeLa cells (Fig. 4G), suggesting that M109S is not a universal cell death inhibitor. Etoposide induces a certain level of cell death even in the absence of Bax and Bak (Han *et al*., 2019; Tu *et al*., 2009). In the case of HeLa cells, Etoposide possibly activated the Bax/Bak-independent pathway inducing apoptosis that cannot be inhibited by M109S. In the case of BAI-1, a previously reported Bax inhibiting compound, it was reported that BAI-1 was not able to inhibit Bax-induced apoptosis of the cancer cells due to the abnormally high levels of Bax expression in the cancer cells(Amgalan *et al*., 2020). Similar to BAI-1, M109S may show selective protection against cell death depending on the expression levels of Bax and other apoptosis regulating proteins.

M109S protected photoreceptor cells from the bright light-induced apoptosis in two mouse models of light-induced retinal degeneration related to Stargardt’s disease and age-related macular degeneration (Maeda *et al*., 2009) (Sawada *et al*., 2014) (Ortega *et al*., 2021). Previous studies showed that Bax plays an essential role in photoreceptor death in these retina degeneration mouse models(Maeda *et al*., 2009) (Sawada *et al*., 2014) (Ortega *et al*., 2021). The photoreceptor protection by M109S confirms that M109S penetrates the blood-retina barrier and functions as Bax inhibitor *in vivo*. Indeed, M109S was detected in the brain and retina in rats 24 h after oral administration (Fig. 2G). Of note, no significant tolerability issues were detected in the mice treated with M109S in these experiments. In addition, no significant off-target activities were noticed at the concentration inhibiting apoptosis (EC50=23.4 nM) in Eurifine’s 78 neurotoxicity tests (please see Supplemental Information of Eurifine Safety Scan test). Although exploratory toxicity studies are planned, the currently available data suggest that M109S may prevent the unwanted death of essential cells without significant side effects.

Bax biding activity of CSMs was confirmed by the MST binding assay. To be noted, the binding of CSMs to Bax was detected in NP40-containing buffer but not in CHAPS-containing buffer (Fig. S5). It is known that NP40, but not CHAPS, induces a conformation change (N-terminus exposure) of Bax which is one of the activation steps of Bax(Hsu and Youle, 1998; Nechushtan *et al*., 1999). Therefore, the present data suggest that CSMs bind partially active Bax (the N-terminus exposed Bax). The Bax binding character of CSMs is similar to that seen in anti-apoptotic Bcl-2 family proteins such as Bcl-2 and Bcl-XL (Hsu and Youle, 1998; Nechushtan *et al*., 1999). For example, Bax binds Bcl-XL only in NP40-based buffer but not in CHAPS-containing (or detergent-free) buffer if cell lysates were prepared from healthy cells (Hsu and Youle, 1998). It is known that Bcl-XL, at the surface of the mitochondrial membrane, interacts with activated Bax (the N-terminus exposed Bax) and bounces Bax back to the cytosol (Edlich et al., 2011) (Todt et al., 2013). As a result, Bcl-XL suppresses the mitochondrial translocation of Bax as well as the accumulation of active Bax in the cell, though the detailed mechanism of this process is not yet understood. Similar to Bcl-XL, CSMs attenuated the N-terminus exposure of Bax and suppressed mitochondrial translocation of Bax. CSMs and Bcl-XL may share a similar mechanism of action to suppress Bax-induced cell death. Further biochemical and structural biological studies are warranted to uncover the precise mechanism of action of Bax inhibition by CSMs.

We found that M109S affects metabolism in 2 ways; 1) by decreasing mitochondrial oxygen consumption and 2) by increasing glycolysis. It has been known that inhibitors of mitochondrial oxidative phosphorylation (OXPHOS) have activities rescuing cells from mitochondria-dependent cell death, especially in the case of ischemia/reperfusion-induced cell death (Gohil et al., 2010; Morciano et al., 2018) (Hong and Pedersen, 2008) (Johnson and Ogbi, 2011) (Grover and Malm, 2008) (Matsuyama et al., 1998). Furthermore, there is increasing evidence showing OXPHOS inhibitors protect neurons in mouse models of neurodegenerative diseases including Alzheimer’s and Parkinson’s diseases (Procaccio *et al*., 2014) (Cunnane *et al*., 2020; Gao *et al*., 2021; Stojakovic *et al*., 2021). These neuroprotective effects of OXPHOS inhibitors have been explained by their activities decreasing ROS by suppressing mitochondrial oxygen consumption (Procaccio *et al*., 2014) (Cunnane *et al*., 2020; Gao *et al*., 2021; Stojakovic *et al*., 2021). As shown in Fig. 6, M109S decreased ROS in MEF cell culture. Therefore, in addition to the direct inhibition of Bax, the suppression of mitochondrial activities may contribute to the cell protection activity of M109S. However, these activities of M109S against mitochondrial activities and metabolism became evident at 1 μM concentration in the cell culture (Fig.6) which is approximately 40 times higher than M109S’s EC50 (23.4 nM) of cell death inhibition (Fig.2). Therefore, the effects on mitochondrial activity may become important only when M109S is used at high dose.

In summary, we developed novel small compounds protecting cells from mitochondria-dependent cell death. The pharmacokinetics of M109S is ideal for *in vivo* treatment. Importantly, M109S reached both the brain and retina, indicating that M109S penetrates the blood-brain/retina barrier. Bax-mediated and mitochondria-initiated cell death are involved in various types of degenerative diseases including neurodegenerative disorders and cardiovascular dysfunctions. M109S and its derivatives have the potential to become an important therapeutic agent that can be used to prevent unwanted cell death in these pathological conditions. The investigation of the mechanism of action of CSMs may shed new light on the previously unknown cell death mechanisms controlled by Bax and Bak.

### Limitation of the study

In the present study, we found that M109S prevented blindness of mouse models of Stargardt disease and macular degeneration. However, these protective effects in mouse disease models do not guarantee that M109S can prevent blindness in human and other animals. We plan to extend our studies in other animals in the near future if fundings were obtained. In the present study, we used a selected group pf cell lines (MEFs, Neruo2a, ARPE19, and HeLa cells) were used to investigate the mechanism of cell death inhibition by M109S. It is not yet certain whether the similar results can be seen in other cell lines. We plan to examine other types of primary cultured cells and cancer cell lines to further investigate the mechanism of action of M109S.

## KEY RESOURCE TABLE

| REAGENT or RESOURCE | SOURCE | IDENTIFIER |
| --- | --- | --- |
| **Antibodies** |  |  |
| anti-Bax polyclonal antibody | ProteinTech | 50599-2-IG |
| anti-Bax Rabbit Polyclonal antibody | BD Pharmingen | BD 554104 |
| Agarose beads conjugated with anti-Bax monoclonal antibody (clone 6A7) | Santa Cruz | sc-23959AC |
| anti-Bak | Santa Cruz | sc-832 |
| anti-Bcl-XL | Santa Cruz | sc-1690 |
| anti-Caspase-3 | Cell Signaling | 9662s |
| anti-beta actin | Sigma-Aldrich | A5441 |
| Anti-rabbit IgG-HRP | Kindle Bioscience | R1006 |
| Anti-mouse IgG-HRP | Kindle Bioscience | R1005 |
| Goat-anti-Mouse IgG (H+L) | Life Tech | G-21040 |
| **Culture medium** |  |  |
| DMEM | ThermoFisher | 11995 |
| Fetal Calf Serum | Fisher Scientific | 12999102 |
| Doxycycline Hyclate | SigmaAldrich | 33429-100MG-R |
| Puromycin | Dot scientific | DSP33020-0.025 |
| Geneticin (G418) | ThermoFisher Scientific | 10131027 |
| Penicillin/Streptomycin | Fisher | SV30010 |
| **Chemicals, peptides, and recombinant proteins** |  |  |
| (2-Hydroxypropyl)-beta-cyclodextrin (bcd) | Sigma-Aldrich | 332607 |
| Etoposide | Sigma-Aldrich | E1383 |
| ABT-737 | Sellekchem | S1002 |
| Obatoclax | Cayman Chemical | 11499 |
| Staurosporine | Sigma-Aldrich | No. S6942 |
| Dexamethasone | Sigma-Aldrich | D4902 |
| M5 (R, S) | This study |  |
| M6 (R, S) | This study |  |
| M7 (R, S) | This study |  |
| M11 | This study |  |
| M109S | This study |  |
| Bax Channle Blocker | Tocris | No. 2160 |
| iMac2 | Tocris | No. 3749 |
| Elthrompag | Sigma-Aldrich | SML3182 |
| **Critical commercial assays** |  |  |
| Capsase-Glo 3/7 Assay kit | Promega | G8093 |
| ROS-Glo-H<sub>2</sub>O<sub>2</sub> Assay kit | Promega | G8820 |
| His-Tag labeling Kit RED-tris-NTA 2<sup>nd</sup> generation | Nanotemper | MO-L018 |
| XF Cell Mito Stress test kit | Agilent | 103015-100 |
| **Deposited data** |  |  |
| NA |  |  |
| **Experimental models: Cell lines** |  |  |
| iWT (wild type MEF, C57BL6 from The Jackson Lab) | This study |  |
| iBBKO (Bax Bak double KO MEFs) | This study |  |
| iRFP (mCherry inducible iBBKO) | This study |  |
| iBax (Bax inducible iBBKO) | This study |  |
| iBak (Bak inducible iBBKO) | This study |  |
| HeLa | ATCC | CRM-CCL-2 |
| Neuro2a | ATCC | CCL-131 |
| ARPE19 | ATCC | CRL-2302 |
| **Experimental models: Organisms/strains** |  |  |
| Balb/c mouse | Jackson Lab | 000651 |
| Abc4<sup>-/-</sup>Rdh8<sup>-/-</sup> mouse | Jackson Lab | 030503 |
| **Recombinant DNA** |  |  |
| pLVX-EF1a-TET3G | Takarabio | 631359 |
| pLVX-TRE3G-mCherry | Takarabio | 631360 |
| **Software and algorithms** |  |  |
| Prism | Dotmatics Graphpad | https://www.graphpad.com/scientific-software/prism/ |

## STAR Methods

### RESOURCE AVAILABILITY

#### Lead contact

Further information and requests for resources and reagents should be directed to and will be fulfilled by the lead contact, Shigemi Matsuyama.

#### Materials availability

There are restrictions on the availability of all newly synthesized chemicals and cell lines used in this study due to potential requirements for materials transfer agreements with the host institution where these materials were generated.

### Data and code availability

Any additional information required to reanalyze the data reported in this paper is available upon request from the lead contact.

## METHOD DETAILS

### Cell Culture and Generation of iRFP, iBax, and iBak Cells

Mouse embryonic fibroblasts (MEFs) from *wt* and *bax^-/-^bak^-/-^*mice (C57BL/6J background, generated by crossing *bax^+/-^* and *bak^-/-^* mice originally obtained from The Jackson Lab) were established in Matsuyama lab, and were immortalized by SV40LT expression. To introduce Tet-ON system (Tets-On inducible system, Takara, Madison, WI), a set of two plasmids were introduced to *bax^-/-^bak^-/-^* MEFs to establish each cell line. The pLVX-EF1a-TET3G carrying Tet-inducible transcription factor was used for all three cell lines. pLVX-TRE3G-mCehrry (original plasmid with only mCherry), pLVX -TRE3G-mCherry carrying cDNAs of human Bax or human Bak, were introduced to generate cell lines of iRFP, iBax, and iBak respectively. These plasmids were introduced into the cells by using Lentivirus transfection system. Doxycycline (Dox) (100 ng/ml) was used to activate Tet-ON system to induce mCherry, Bax, and Bak. To maintain transfected genes, puromycin (1 ug/ml) sand G418 (750 ug/ml) were added to the MEF medium. The MEF medium is made from DMEM High Glucose (ThermoFisher Catlog Number11995) supplemented with 10% Fetal Calf Serum (FCS, Fisger Scientific Cat No.12999102), 2 mM (final concentration) Sodium Pyruvate, 6 mM L-Glutamine (Final concentration), 100U/ml (Penicillin-Streptomycin), and 1% of Non-Essential Amino Acid supplement (Gibco, 11140-050).

### High-Throughput Screening of Small Compound Library

We used a 50,000 small compound library purchased from ChemBridge (San Diego, CA) to identify the compounds protecting cells from Bax-induced cell death using iBax cells (passage of iBax cells were used in the screening). 386 well plates and Operetta Image analysis system (Perkin Elmer) were used for the screening. First, each compound was added together with Dox (100 ng/ml) to the culture media of iBax cells to the final concentration of 10 µM. Cells were cultured for 48 h and fixed by 2% paraformaldehyde (PFA) and the nuclei were stained with Hoechst 33258. Cell images were captured using both RFP and DAPI filter channels. RFP filter channel was used to monitor the mCherry expression. The compounds maintaining 80% and higher mCherry expression of the control cells (only Dox addition) were analyzed. DAPI filter channel was used to analyze the percentage (%) of cells with apoptotic nuclei (pyknosis). The best 50 chemicals were selected, and they were subjected to examine the dose-dependent effects to confirm the cytoprotective activities. After the best 50 hit compounds were selected, and two groups of chemicals were found Group 1 (M5 and M6 were the best Bax inhibitors in this group) and Group 2 (M7 was the best Bax inhibitor in this group)). We examined whether these 50 chemicals can also inhibit Bak-induced cell death in iBak cells. However, these compounds were not effective in inhibiting Bak-induced cell death in iBak cells.

### Chemical Synthesis and profiling tests

All the chemical synthesis were performed by Wuxi AppTech (Hong-Kong, China) based on the chemical design by Dr. William Greenlee. To select the lead compounds, the hit compounds were subjected to the profiling tests examining, kinetic solubility, Caco-2 cell, microsomal stability (t ½ min), and plasma protein binding.

### Western Blot

To analyze the expression levels of Bcl-2 family proteins, Caspase-3, and beta-tubulin, cells were lysed in RIPA buffer and 20 micro-g of total protein was subjected to Western blot analysis using 4-20% gradient gel (Invitrogen). The primary antibodies used in this study are: anti-Bax antibody (Proteintech, 50599-2-IG, 5000X), anti-Bcl-2 antibody (SantaCruz, sc-7382, 500X), anti-Bc-XL antibody (SantaCruz, sc-1690, 500X), Caspase-3 (Cell Signaling, 9662, 1000X), and anti-beta actin (Sigma Aldrich, A5441, 20,000X). The secondary antibodies used for this study were anti-Rabbit-HRP (Kindle Biosciences, R1006, 1000X) and anti-mouse IgG-HRP (Kindle Biosciences, R1005, 1000X). Western blot signals were captured by Kwik Quant Imager (Kindle Biosciences), and the densitometric analysis was performed by Image J.

### Caspase Measurement

Promega’s Capsase-Glo 3/7 Assay kit (G8093, Promega) was used to measure caspase activity according to the manufacturer’s protocol. i<u>Bax and MEFs (ABT-737-and Staurosporine (STS)-</u> <u>induced apoptosis)</u>: iBax cells or MEFs were seeded at the density of 16,000 cells/100 μl/well (for ABT-737) or 20,000 cells/100 μl/well (for STS) in 96 well plates on Day 0. For culture medium, MEF medium was used (please see “Cell Culture” section). To determine the effects of CSMs on Caspase activation by Bax overexpression, Dox (100 ng/ml) and CSMs (various concentrations as indicated in the figures) were added to the culture media on Day 1. On Day 3, cells were subjected to Caspase assay. To examine the effects of CSMs against 1 µM ABT-737 (S1002, Sellekchem), cells were treated with ABT-737 together with various concentrations of CSMs. On Day 1 (*wt* (passage 10) and Bax only (passage 11) cells) and Day 2 (Bak only cells (passage 10), Bak only cells die slower than wt and Bax-only cells)), Caspase activity was measured. To examine the effects of M109S against STS-induced apoptosis, MEFs were pre-incubated with M109S for 2 h before STS (1 µM) addition to the medium. Four hours after STS addition, Caspase assay buffer containing cell lysis reagent was added to each well, and cells were subjected to Caspase measurement. <u>Neuro2a and HeLa cells (Etoposide-induced apoptosis:</u> Cells were seeded at the density of 10,000 cells/100 μl/well in 96 well plates on Day 0. On day 1, etoposide (25 µM to Neuro2a (ATCC, passage 12), 12.5 or 25.0 µM or HeLa cells (ATCC, passage 11), respectively) were added to the culture media. On day 2, media were replaced with fresh media containing various concentrations of M109S without etoposide. On day 3, Caspase assay buffer was added to each well and Caspase activity was measured.

### Pharmacokinetic Analysis

The compounds were dissolved in 15% (2-Hydroxypropyl)-beta-cyclodexitrin (bCD) (and they were administered to the mice or rats by oral gavage, intrapreneurial (i.p.), or intravenous (i.v.) injection. At the time point of 0.5, 1, 2, 4, 8, and 24 h, the concentration in the blood plasma (3 mice for each time point) were measured by Mass Spectrometry analysis. The concentrations in the retina and brain were also measured 24 h after the administration. These PK analyses of mice and rats were performed by Wuxi AppTech (Hong-Kong, China) and PharmOptima (Portage, Michigan).

#### Animals Care and Treatment

Double knockout mice *Abca4^−/−^Rdh8^−/−^* with 129Sv or C57BL/6 background that lacks ABCA4 transporter and retinol dehydrogenase 8 (RDH8) (Maeda et al., 2008) at 5–6 weeks of age were used to evaluate the protective effect of a novel BAX inhibitor (M109S) on retinal degeneration induced by acute light. *Abca4^−/−^Rdh8^−/−^*mice do not carry the *Rd8* mutation, however, they carry the Leu variation at amino acid 450 of retinal pigment epithelium 65 kDa protein (RPE65) (Gao et al., 2018; Kim et al., 2004). The substitution of Leu to Met in the *Rpe65* gene that exists in inbreed mice decreases sensitivity to retinal damage induced by bright light (Kim *et al*., 2004). Balb/cJ (RRID:IMSR_JAX:000651) mice were used as WT control. Both male and female mice were used in all experiments. All mice were housed in the Animal Resource Center at the School of Medicine, Case Western Reserve University (CWRU), and maintained in a 12-hour light/dark cycle. All animal procedures and experimental protocols were approved by the Institutional Animal Care and Use Committee at CWRU and conformed to recommendations of both the American Veterinary Medical Association Panel on Euthanasia and the Association for Research in Vision and Ophthalmology.

#### Retinal Degeneration Induced with Bright Light

Twenty four hours before the treatment 4-5 weeks old *Abca4^−/−^Rdh8^−/−^*mice and 6-8 weeks old Balb/cJ mice were dark-adapted. M109 was dissolved in the 2-hydroxypropyl-β-cyclodextrin vehicle and administered to mice by gavage. *Abca4^−/−^Rdh8^−/−^* mice were administered with M109S at 1, 2.5, 5, 10, and 20 mg/kg body weight (b.w.) 24 h and 2 h before the exposure to light, while Balb/cJ mice were gavaged with M109S at a dose of 20 mg/kg b.w. 2-hydroxypropyl-β-cyclodextrin vehicle was administered to control mice. To initiate the retinal damage, mice pupils were dilated with 1% tropicamide and *Abca4^−/−^Rdh8^−/−^*mice were exposed to 10,000 lux white light for 30 min, delivered from a 150-W bulb (Hampton Bay; Home Depot, Atlanta, GA) (Chen et al., 2012). Balb/cJ mice were exposed to 12,000 lux light for 2 h. After illumination, mice were kept in the dark for 7 days. Treatment with M109 or vehicle was repeated 24 and 48 h after the exposure to light. The health of the retina was assessed *in vivo* by imaging using spectral domain optical coherence tomography (SD-OCT). Next, mice were euthanized and eyes were collected for histological evaluation. Six mice were used in each experimental group.

#### SLO

The Scanning Laser Ophthalmoscopy (SLO) (Heidelberg Engineering, Franklin, MA) was used to image the *in vivo* whole-fundus mouse retinas. *Abca4^−/−^Rdh8^−/−^*mice were subjected to SLO imaging immediately after the acquired SD-OCT imaging. The number of autofluorescent spots (AF) detected in different experimental groups was quantified and compared to determine the statistical significance. Six mice were used in each experimental group.

#### SD-OCT

Ultrahigh-resolution SD-OCT (Bioptigen, Morrisville, NC) *in vivo* imaging was used to evaluate the effect of M109S treatment on retinal structure in *Abca4^−/−^Rdh8^−/−^* mice subjected to bright light-induced retinal damage (Chen *et al*., 2012). Before imaging, mice pupils were dilated with 1% tropicamide and anesthetized by i.p. injection of a cocktail containing ketamine (20 mg/ml) and xylazine (1.75 mg/ml) at a dose of 4 µl/g b.w. The a-scan/b-scan ratio was set at 1200 lines. The retinal images were obtained by scanning at 0 and 90 degrees in the b-mode. Five image frames were captured and averaged. To determine the changes in the retinas of mice treated with M109S and exposed to bright light and control mice treated with vehicle or non-treated and dark-adapted mice were determined by measuring the outer nuclear layer (ONL) thickness at 0.15-0.75 mm from the optic nerve head (ONH). Values of the ONL thickness were plotted using means and standard deviation. For each experimental group, six mice were used.

#### Retinal Histology

The effect of M109S on the retinal morphology in *Abca4^−/−^Rdh8^−/−^*mice exposed to bright light was determined by retinal histology analysis. Eyes were collected from non-treated mice kept in the dark, treated with 2-hydroxypropyl-β-cyclodextrin vehicle or M109S, and illuminated with bright light. Eyes were collected from euthanized mice and fixed in 10% formalin in PBS for 24 h at room temperature (RT) on a rocking platform, followed by paraffin sectioning. Sections (5 µm thick) were stained with hematoxylin and eosin (H&E) and imaged by a BX60 upright microscope (Olympus, Tokyo, Japan). Then, the data were processed using MetaMorph software (Molecular Devices, Sunnyvale, CA, USA). Six mice were used in each experimental group.

#### Immunohistochemical Analysis

To detect the rod and cone photoreceptors in the retina, eyes were collected from euthanized *Abca4^−/−^Rdh8^−/−^* mice one week after the treatment and fixed in 4% PFA for 24 h followed by their incubation in 1% PFA for 48 h at RT. The eight-μm thick cryosections prepared from fixed eyes were incubated at 37°C for 10 min, followed by hydration with PBS for 2 min. These sections were blocked with 10% normal goat serum (NGS) and 0.3% Triton X-100 in PBS for 1 h at room temperature. Then sections were stained with a mouse 1D4 anti-rhodopsin primary antibody to visualize rods and biotinylated peanut agglutinin (PNA) (1:500 dilution) to visualize cones overnight at 4°C. The next day, sections were washed with PBS, followed by the incubation with Alexa Fluor 555-conjugated goat anti-mouse secondary antibody (1:400 dilution) to detect rods and Fluor 488-conjugated streptavidin (1:500 dilution) to detect cones for 2.5 h at room temperature. Cell nuclei were stained with DAPI. Slides were coverslipped with Fluoromount-G (SouthernBiotech). The retina was imaged with the Olympus FV1200 Laser Scanning Microscope (Olympus America, Waltham, MA) and 40x/1.4 objective.

### 6A7 Ab Bax Immunoprecipitation

Immunoprecipitation of activated Bax (The N-terminus exposed Bax) was performed as previously reported (Gama et al., 2009). Cell lysates were prepared with 1% CHAPS buffer. The cell lysates of 150 ul (5 mg/ml protein concentration) were incubated with 10 ul of 6A7 antibody-conjugated agarose beads (sc-23959AC, Santa Cruze) at 4C for 2 hrs. Then, the beads were washed with the CHAPS buffer 3 times, and the immunoprecipitated Bax was eluted by incubating the beads with 50 ul of Lamili buffer at 95C for 10 min. Fifteen (15) ul of the eluted samples was subjected to Western blot. For the detection of the total Bax expression in the cell lysate, 20 μg of total protein was subjected for each lane of the gel. To detect Bax in Western blot, an anti-Bax polyclonal antibody (50599-2-IG, Protein Tech) was used.

### Bax Immunocytochemistry

On Day 0, iBax cells (15,000 cells/100 μl/well) were seeded in 96 well plates, On Day 1, Dox (100 ng/ml) was added together with CSMs. After 36 h of Dox addition, cells were fixed by 2% PFA and subjected to immunocytochemistry. Anti-Bax polyclonal antibody (50599-2-IG, Protein Tech) was used to detect Bax in the cells. The pictures of Bax staining (green fluorescence), mCherry expression (red fluorescence), and normal light image were taken and used to determine the percentages of cells showing punctuated Bax staining among Bax expressing cells. At least 200 cells were counted from each image. At least 3 wells were used for each condition.

### Microscale Thermophoresis (MST)

His-Tag labeling Kit RED-tris-NTA 2^nd^ generation (MO-L018, Nanotemper) was used to label His-tagged Bax purchased from Creative BioMart (Catalog No. BAX-6976H, Seattle, WA). His-Bax (210 µl/ml in Phosphate Buffer Saline (PBS) (pH7.4)) was aliquoted (10 µl/tube) and kept at -80C. For RED labeling, 0.05% CHAPS PBS was used to dissolve the RED dye. The final concentration of the labeled Bax protein was 50 nM for the binding assay and was mixed with a serial dilution of CSMs. The first concentration of CSMs (20 µM) was prepared in PBS with 1% BSA and 2% NP40. The final condition of the buffer for binding was PBS with 0.5% BSA and 1% NP40 in all 16 reaction tubes. When 1% NP40 was replaced with 1% CHAPS, the dose-dependent binding signal was not detected as explained in Discussion. The labeled His-Bax was kept on ice and used within 3 hrs after the labeling. The measurement of dissociation constant Kd was performed by the NanoTemper’s software installed in the Monolith system.

### Seahorse Experiment

XF Cell Mito Stress test kit (103015-100) was used to measure OCR and ECAR with the XFe96 Seahorse instrument. On Day 0, the cell culture of MEFs was started (1.6×10^4^ cells/200 micro-l/well (Agilent, XF96 microplate V3-PS)) using a MEF medium. On Day 1, CSMs were added to the culture medium (10 nM, 100nM, or 1 µM) and incubated for 4 hrs in a CO_2_ incubator. After the incubation, the medium was changed with XF DMEM medium (Agilent, 103575-100), and cells were incubated in a 37 °C chamber (100% air) for 1 hr before the mitochondria stress test. For the test, rotenone (0.5 µM), antimycin A (0.5 µM), oligomycin (1.5 µM), and FCCP (2.0 µM) were used according to the manufacturer’s protocol. OCR and ECAR were recorded and the effects of CSMs on OXPHOS and glycolysis were analyzed.

### ROS Measurement

ROS-Glo-H_2_O_2_ Assay kit (Promega G8820) was used to measure ROS in MEFs culture (medium and cells) according to the manufacturer’s protocol. On day 0, cells were seeded at the concentration of 16,000 cells/100 μl/well in 96-well plate. On day 1 (24 hr after the start of cell culture), cells were pre-incubated with DMSO only (Control) (0.001%) or M109S (100 nM, 200 nM, 1 µM (the final DMSO was adjusted to 0.001% in all the conditions) for 4 hrs. Then, the medium was changed to a new medium containing 25 µM H2O2 substrate with DMSO or M109S (medium in each well was 100 μl). Cells were cultured in a CO_2_ incubator for additional 4 hrs. Then, 100 μl of ROS-glo Detection solution was added to each well and incubated for 20 min at room temperature. Luminescence intensity from each well was recorded by MG3500 (Promega) plate reader.

### Cell Number Measurement

On Day 0, the cell culture of MEFs was seeded (4×10^5^ cells/ml/well (12 well plate)). On day 1, 24 hrs after the starting of the cell culture, the various concentration of CSMs were added to the culture medium. On Day 2, 48 hrs after the start, cells were collected, and cell number was counted by an automated cell counter (Countess II, Invitrogen). Four wells were used for one condition.

### Statistical Analysis

For Fig.1, 2, 4, 5, and 6, Statistical analyses and calculations of EC_50_ were performed using Prism (version 9.4.1). Student’s t-test was used to determine the statistical significance in *in vitro* experiments. For Fig.3: The statistical analysis of treatment with M109S in mouse models one or two-way ANOVA with Turkey’s post hoc tests were used. All statistical calculations were performed using the Prism GraphPad 7.02 software. Type 1 error tolerance for the experiments was assumed at 5%. Values of *P* < 0.05 were considered statistically significant. Data collection and statistical analysis were performed by different personnel.

## Author Contribution

Shigemi Matsuyama initiated the project and designed the experiments reported in Fig. 1, 2, 4, and 5. iRFP cells, iBax cells, and iBak cells as well as the CSM screening system were designed and generated by Shigemi Matsuyama. Mieko Matsuyama performed the experiments together with Shigemi Matsuyama reported in Fig. 1, 2 (except Fig. 2 E-G), 4, and 5. Jonah Scott-McKean analyzed the data of Fig. 1F. Joseph Ortega and Beata Jastrzebska designed and performed the experiments reported in Fig. 3. Yuri Fedorov and Drew Adams set up the high-throughput drug screening system with the Operetta system (Parkin Elmer), optimized the CSM screening system, and performed the screening. Jeannie Muller-Greven and Mathias Buck provide advice for the MST-binding assay and participated in the data analysis. William Greenlee performed the medicinal chemistry study during the entire project, and he designed all the novel CSMs. The first draft of the manuscript was prepared by Shigemi Matsuyama, and all the authors participated in editing the draft to complete the final version.

## Acknowledgement

Authors are grateful for invaluable advice and guidance by Dr. Diana Wetmore (HDI) and Dr. Steven Brenner (HDI) throughout the project, and for project management by Dr. Jeff Klein (HDI). We thank the past and present members of Matsuyama laboratory (Dr. James Palmer, Dr. Kelsey Jensen, David J WuWong, Sean Wang, Emma Heironimus, and Xiao-Yi Chen) for their assistance in plasmid production, cell image capture, and data analysis. We also thank Dr. David Wald for his advice on drug screening and the use of the Tet-ON system. This work was supported by Gund-Harrington Scholar Award (to SM) from Foundation Fighting Blindness and Harrington Discovery Institute (HDI), Department of Defense Vision Research Program W81XWH-20-1-0735 (to SM), National Institute of Health (NIH) RO1-AG031903 (to SM), RO1 EY032874 (to BJ), RO1EY029169 (JMG and MB), P30 grants for Case Comprehensive Cancer Center and Vision Science Research Center.

---

**Supplemental Figure S1**

The effect of M109S concentration on retina inflammation was examined *in vivo* in the whole fundus images of *Abca4^-/-^Rdh8^-/-^* mice taken with scanning laser ophthalmoscopy (SLO) one week after exposure to bright light. Autofluorescence spots (AF) representing macrophages and microglial cells activated by light injury to clear dying photoreceptors were detected in vehicle-treated mice, but not in M109S-treated mice at 10 mg/kg/b.w. However, lower doses of M109S were not effective. Scale bar, 1 mm.

**Supplemental Figure S2**

Concentration-dependent effect of M109S protecting against light-induced retina degeneration. The Spectral Domain Optical Coherence Tomography (SD-OCT) images of the retina from *Abca4^-/-^Rdh8^-/-^* mice were taken one week after illumination. Scale bar, 100 µm. The exposure to bright light-induced cell death of photoreceptors, which caused shortening of the outer nuclear layer (ONL) in the retinas of vehicle-treated control mice. Retina degeneration was prevented by M109S at 10 and 20 mg/kg/b.w. However, lower doses (1, 2.5, and 5 mg/kg) were not effective.

**Supplemental Figure S3**

M109S at 10 mg/kg b.w. protected photoreceptors of Balb/cJ mice from light-induced cell death. As detected in the OCT images (A) and H&E stained the retina sections (B) the thickness of the ONL layer resembled the one observed in dark-adapted control mice, while vehicle-treated mice showed severe shortening of the ONL as a result of photoreceptor cells death. Scale bar in (A) 100 µm and in (B) 50 µm.

**Supplemental Figure S4**

M109S was not able to inhibit etoposide-induced apoptosis in HeLa cells. HeLa cells were treated with etoposide (25 µM) for 24 h. Then, the medium was changed, and cells were incubated with various concentrations of M109S for an additional 24 h without etoposide. M109S did not show a significant inhibition of etoposide-induced caspase activation in HeLa cells, except for a slight inhibition at 125 nM. The result of the Student’s t-test statistical analysis is shown in each graph. “ns”: No statistically significant difference was detected (p>0.05).

**Supplemental Figure S5**

Microscale thermal shift (MST) assay detecting the binding of CSMs and Bax. A: The binding of Bax and M41S was detected when the interaction was determined using 1% NP40-containing buffer. B: The binding of Bax and M41S was not detected in 1% Chaps buffer.

