## Supplemental Figures for "Development of novel cytoprotective small compounds inhibiting mitochondria-dependent apoptosis"

Supplemental Figure 1

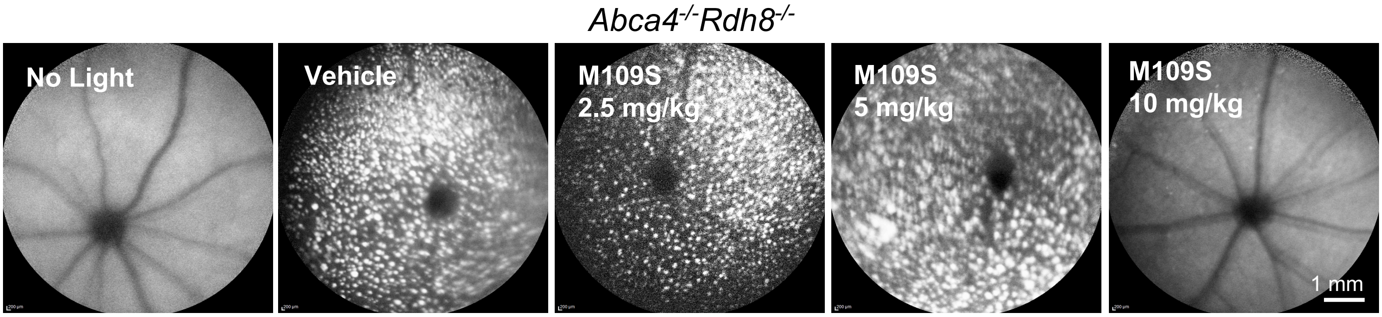

### Supplemental Figure 2

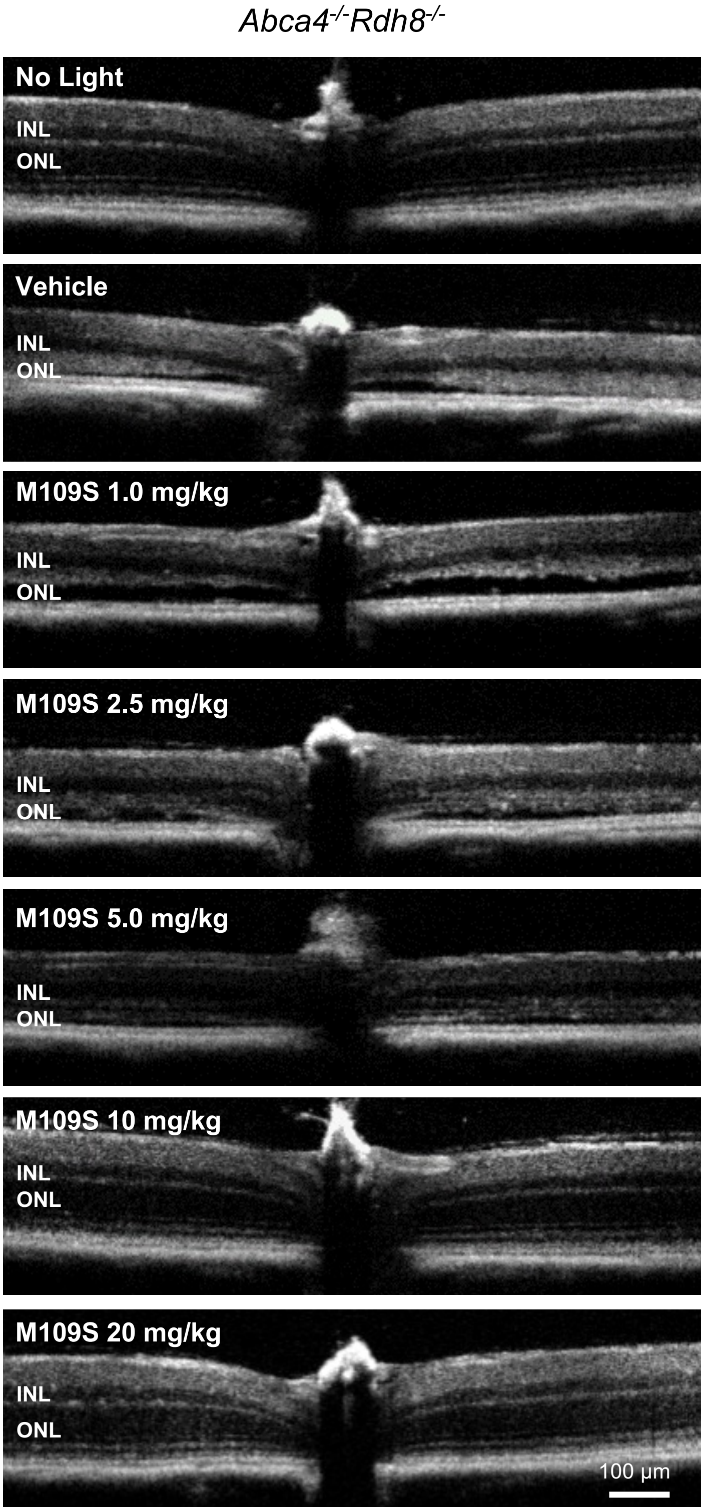

### Supplemental Figure 3

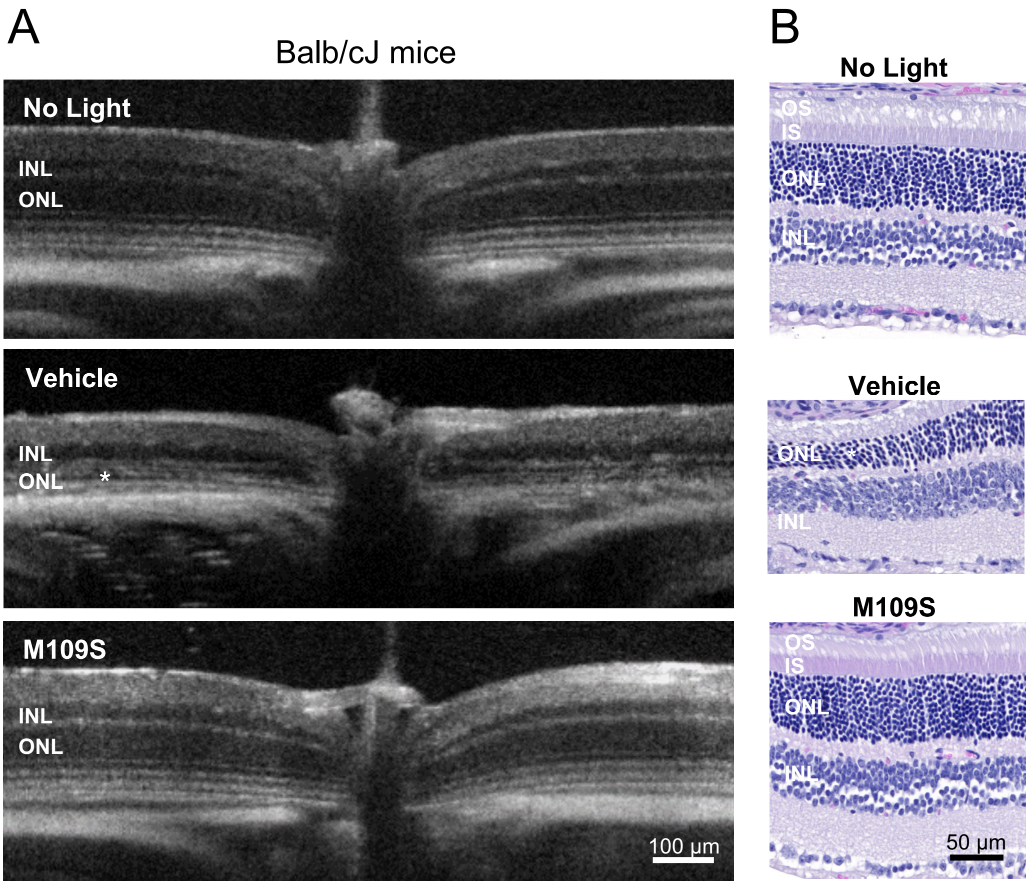

Etoposide (25 uM)-induced apoptosis in HeLa cells:
M109 did not show a significant apoptosis inhibition.

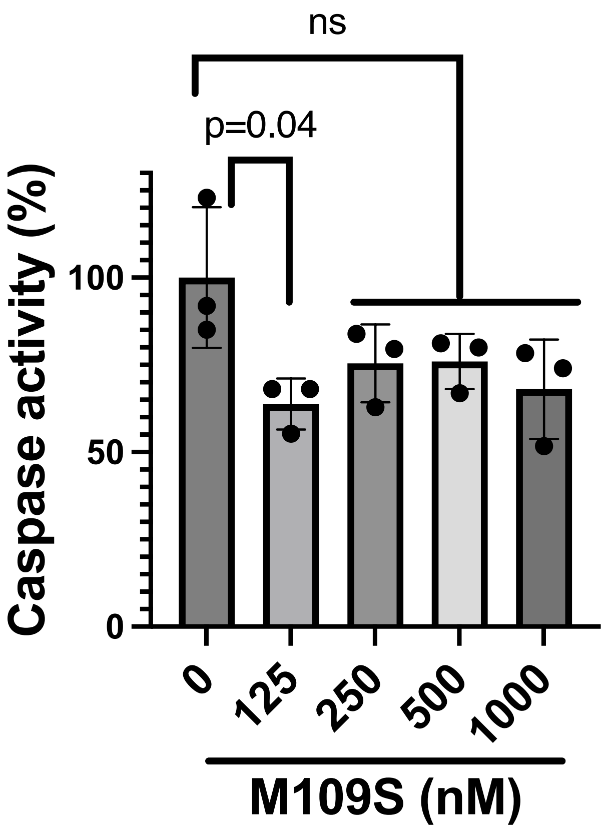

### Supplemental Figure 5

A

**Binding analysis in NP40 Buffer**

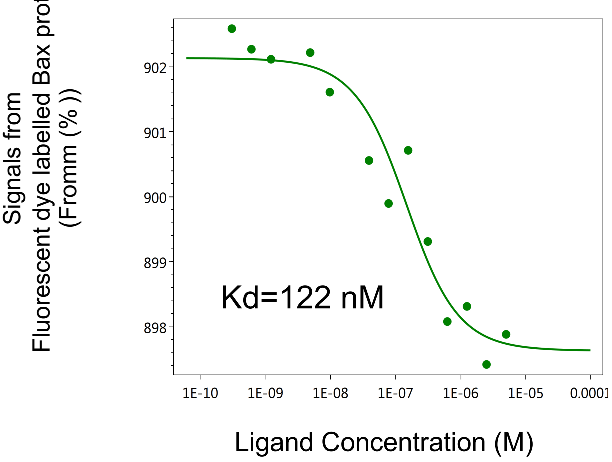


B

**Binding analysis in CHAPS Buffer**

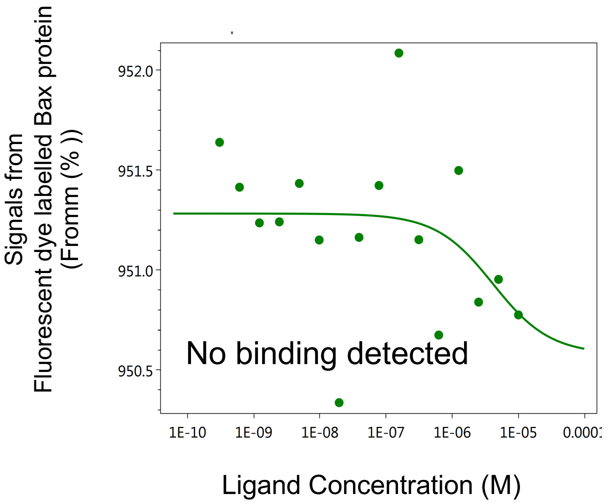
